# ontophylo: Reconstructing the evolutionary dynamics of phenomes using new ontology-informed phylogenetic methods

**DOI:** 10.1101/2023.06.13.544734

**Authors:** Diego S. Porto, Josef Uyeda, István Mikó, Sergei Tarasov

## Abstract

1. Reconstructing ancestral states for discrete characters is essential for understanding trait evolution in organisms. However, most existing methods are limited to individual characters and often overlook the hierarchical and interactive nature of traits. Recent advances in phylogenetics now offer the possibility of integrating knowledge from anatomy ontologies to reconstruct multiple discrete character histories. However, practical applications that fully harness the potential of these new approaches are still lacking.

2. This paper introduces *ontophylo*, an R package that extends the PARAMO pipeline to address these limitations. *Ontophylo* enables the reconstruction of phenotypic entities composed of amalgamated characters, such as entire phenomes or anatomical regions. It offers three new applications: (1) reconstructing evolutionary rates of amalgamated characters using phylogenetic non-homogeneous Poisson process (pNHPP), allowing for rate variation and shifts over time and phylogeny; (2) explicit reconstruction of morphospace dynamics; and (3) direct visualization of evolutionary rates and statistics on vector images of organisms. *Ontophylo* incorporates ontological knowledge to facilitate these applications.

3. Benchmarking confirms the accuracy of pNHPP in estimating character rates under different evolutionary scenarios, and example applications demonstrate the utility of *ontophylo* in studying morphological evolution in Hymenoptera using simulated data.

4. *Ontophylo* is easily integrated with other ontology-oriented and general-purpose R packages and offers new opportunities to examine morphological evolution on a phenomic level using new and legacy data.

## 1 INTRODUCTION

Reconstruction of ancestral states for discrete characters is commonly used to understand trait evolution in organisms. Most methods for ancestral reconstruction were developed for individual characters which represent some elementary phenotypic observations with a limited number of states. When we focus on individual traits, phenotypic evolution becomes oversimplified, as organisms exhibit complex systems where traits are arranged hierarchically and interact over time (Rasskin-Gutman and Esteve-Altava, 2014; Esteve-Altava, 2017).

Recent advancements in statistical phylogenetics provide a path toward a solution to this problem. The PARAMO pipeline (Tarasov et al., 2019) allows the reconstruction of phenotypic entities comprising collections of individual traits, including entire phenomes, anatomical regions, or morphological complexes. In essence, it is based on the principle that Markov models, commonly employed to model discrete traits, can be used to construct a composite character from individual ones in a mathematically consistent manner (Tarasov, 2019, 2020). Following this principle, individual characters can be combined by amalgamating their rate matrices. The problem, however, is that such amalgamations produce enormous matrices that are non-computable. As an alternative to rate matrices, the PARAMO approach introduced computationally feasible amalgamation based on stochastic maps (SMs) (Huelsenbeck et al., 2003).

Researchers often seek to understand the mode, tempo, and variation of character rate evolution across a phylogenetic tree, rather than solely focusing on ancestral state reconstruction based on SMs as provided by PARAMO. To address this, we introduce a new R package, called ontophylo, which implements and expands the PARAMO pipeline. `Ontophylo` offers three new applications. (1) A phylogenetic non-homogeneous Poisson process (pNHPP) method for reconstructing evolutionary rates of amalgamated characters based on SMs. This method allows for the inference of rate variation and shifts over both time and phylogeny. (2) The explicit reconstruction of morphospace dynamics for anatomical regions or entire phenomes using SMs and pNHPP. Additionally, `Ontophylo` provides functionality for animating the evolution of reconstructed morphospaces over time. (3) Direct visualization of evolutionary rates and other statistics associated with anatomical regions on organism’s vector images.

For all three applications, `Ontophylo` makes use of ontological knowledge to facilitate and inform the amalgamation of SMs and the mapping of rates to images. Ontologies are structured controlled vocabularies used to describe knowledge in a given domain (Balhoff et al., 2010; Dahdul et al., 2010). In this context, knowledge is represented as a hierarchical graph with formally defined concepts and relationships (Vogt, 2019; Thessen et al., 2020). When a phylogenetic dataset is annotated with ontology terms, researchers can easily query all characters annotated with relevant terms of interest. `Ontophylo` leverages these annotations to automatically construct amalgamated characters for any level of the anatomical hierarchy. Furthermore, `Ontophylo` is capable of visualizing reconstructed rates by using a vector image containing annotated body regions. While using ontologies is recommended, it is not mandatory as `Ontophylo` allows users to provide a custom graph of anatomical hierarchy.

Finally, we discuss how `Ontophylo` can be used to enhance phenomic-scale research in biology and address various questions concerning trait dynamics.

## 2 MATERIALS AND METHODS

### 2.1 Package overview

#### >Ontophylo: PARAMO pipeline

`Ontophylo` implements the PARAMO pipeline (Tarasov et al., 2019) in R. In summary, the pipeline workflow has the following steps.

**Step 1**. As an input, the PARAMO takes a dated phylogeny, a phylogenetic character matrix, and character annotations to anatomy ontology terms (Figure 1a). Phylogenetic characters can be annotated manually with ontology terms using a CSV table or semi-automatically with ontoFAST (Tarasov et al., 2022). It is crucial to ensure the independence of all phylogenetic characters before sampling character histories, as discussed in Tarasov (2019). However, anatomical dependencies are common in most morphological data sets. For example, one character may describe the absence/presence, while another character describes the quality (e.g. shape, color) of an anatomical entity. In such cases, dependent characters should be modeled together and recoded accordingly. `Ontophylo` does not recode characters automatically, but this can be achieved using rphenoscate (Porto et al., 2023).

**FIGURE 1.**
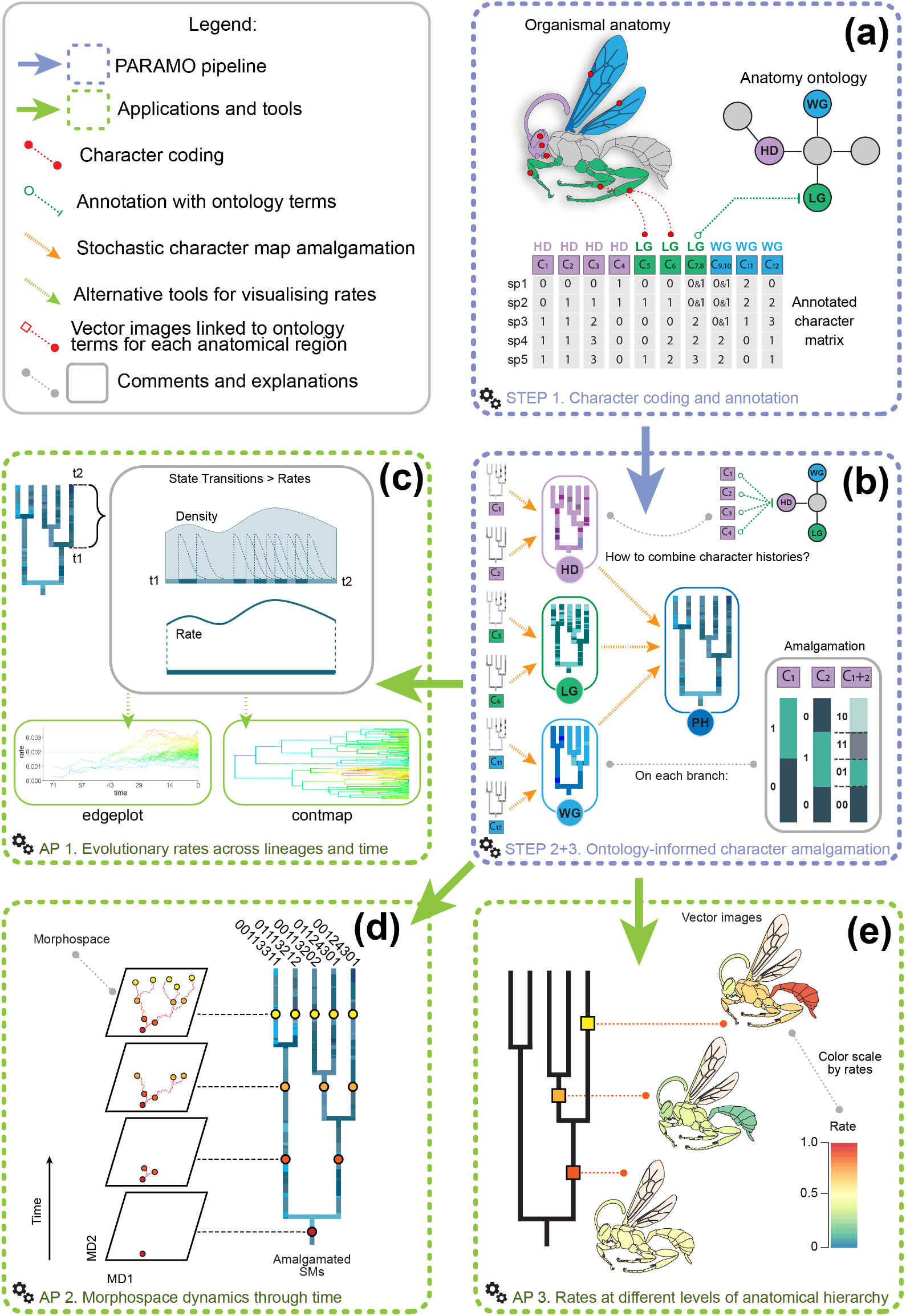
Steps of the PARAMO pipeline and `Ontophylo` applications. **(a)** Character coding and annotation of phylogenetic characters with ontology terms. **(b)** Ontology-informed amalgamation of stochastic character histories. **(c)** Application 1: Estimating rates of character evolution across lineages and time. **(d)** Application 2: Reconstructing morphospace dynamics through time. **(e)** Application 3: Estimating evolutionary rates for morphological complexes, anatomical regions, or phenome. Abbreviations: C1-12, characters; HD, head; LG, leg; PH, phenome; WG, wing.

**Step 2**: In this step, Markov models of discrete character evolution are fitted to each individual character. `Ontophylo` can assist in managing the data for this analysis. The model fitting can be performed using various software, such as corHMM (Beaulieu et al., 2013) or phytools (Revell, 2012) for maximum likelihood estimation, or RevBayes for Bayesian inference (Höhna et al., 2016). It is recommended that users fit multiple alternative models (e.g., equal rates, asymmetric, all rated differently) to the same character and choose the best model using model selection.

**Step 3**: In this step, PARAMO uses stochastic mapping to sample character histories for each individual character. The SM represents the mapped evolutionary history of a character, which is conditioned on the data at the tips and the chosen Markov model (Huelsenbeck et al., 2003). The same software as in the previous step (e.g., corHMM, phytools, or RevBayes) can be used for this purpose. Next, the individual characters are amalgamated based on morphological complexes, anatomical regions, or the entire phenome, following the hierarchy described by the reference anatomy ontology (Figure 1b). The SM amalgamation involves constructing a joint stochastic map by combining time segments from individual SMs. The amalgamation of rate matrices and SMs is equivalent (Tarasov et al., 2019). It is important to note that PARAMO assumes the independence of individual characters during amalgamation, while the dependent evolution is modeled in Step 1. The level of amalgamation is determined by the user and is based on the granularity of primary ontology annotations, which can range from general to specific anatomical terms.

Once amalgamated stochastic character maps were obtained, three applications, summarized below, are possible with `Ontophylo`.

#### >Ontophylo: Reconstructing rates using pNHPP

In this application, `Ontophylo` uses pNHPP to estimate the evolutionary rate of the amalgamated character, allowing for variation across and within tree branches. This rate represents the speed of change between character states. The pNHPP method takes the state change events from the sample of stochastic character maps (SMs) as input and converts them into a reconstructed rate function using a non-parametric approach (see, the technical details below). The pNHPP method relies on minimal initial assumptions regarding the distribution of rates across the phylogeny.

The main one is that the rate represents a continuous smooth function, ensuring smooth transitions for all rate shifts. The variation of the rate over the tree is inferred from the data. The reconstructed rate can be visualized on the phylogeny using the contmap function from phytools or as an edgeplot of rates through time using `Ontophylo`’s edgeplot function (Figure 1c).

#### >Ontophylo: Reconstructing Morphospace dynamics

An amalgamated character can potentially have a large number of states, as it is formed from all possible combinations of states in the initial characters. For example, two binary characters yield an amalgamation with four states: {00, 01, 10, 11}. The similarity between these amalgamated states can be measured using the Hamming distance, which counts the number of positions at which two state strings differ. A SM explicitly reconstructs the evolutionary sequences of ancestral states at each point on the timetree. This information on the reconstructed state sequences and their similarity can be utilized to reconstruct the morphospace. `Ontophylo` offers the functionality to measure Hamming distances between all reconstructed states and states at the tips for a SM. Additionally, the similarity between them can be visualized as a 2D plot using dimensionality reduction techniques (e.g., multidimensional scaling, UMAP). This provides a fine-scale reconstruction of the morphospace. Furthermore, `Ontophylo` can incorporate the time axis of the process by integrating the pNHPP rate with the morphospace. This integration allows for the creation of an animation that demonstrates the evolution of the morphospace over time (Figure 1d).

#### >Ontophylo: Visualizing rates on organismal images

When the pNHPP rate is reconstructed for multiple anatomical regions, it becomes valuable to visualize how it varies across both the phylogeny and body regions. To achieve this, `Ontophylo` offers the ability to link vector images representing organismal body regions with ontology terms and visualize reconstructed rate values on them. As these values can differ among branches, `Ontophylo` supports querying values from specific branches or time points on tree and plotting them on the associated image (Figure 1e).

Besides the overall transition rate, any customizable statistics of interest can be visualized on the body regions. The images used should be in vector format, with distinct graphical objects representing different body regions. These images can be created using software like Adobe Illustrator or GIMP and then imported into R. `Ontophylo` accepts image annotations in the form of a CSV file.

### 2.2 Phylogenetic Non-Homogeneous Poisson Process (pNHPP)

The pNHPP approach is based on the concept of a time-dependent Poisson process, where a collection of state changes in space occurs at a rate *λ*. The number of state changes *N*_*s*_ within a given time interval follows a Poisson distribution *N*_*s*_ ∼ Poisson(*λ* (*t*)). Previous studies (Taddy and Kottas, 2012; Ross and Markwick, 2018) have demonstrated that *λ* (*t*) can be estimated without assuming a specific functional form by decomposing it into an amplitude *λ*_0_, which controls the total number of events, and a probability density function (pdf) *G* (*t*), which determines the distribution of events within the observation window. This results in the equation *λ* (*t*) = *λ*_0_*G* (*t*), where *G* (*t*) satisfies the pdf condition ∫*T G* (*t*)*d t* = 1.

We adopt this fundamental principle to characterize the pNHPP. Consider a phylogenetic timetree that spans the time interval [*T*, 0] with a given number of branches *N*_*b*_ . Each branch is denoted as *t*_*i*_, representing a specific time point (*t*) located on the *i* th branch. Thus, the process rate can vary across branches and within branch segments as *λ* (*t*_*i*_). Consequently, the number of state change events (*N*_*s*_) within a branch segment in a stochastic map is *N*_*s*_ ∼ Poisson(*λ* (*t*_*i*_)). Our goal is to infer *λ* (*t*_*i*_) based on the observed state change events in a sample of SMs.

With the aforementioned decomposition, *λ* (*t*_*i*_) can be expressed as:

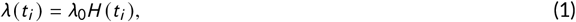

where *H* (*t*_*i*_) represents the pdf defined over the entire timetree. The pdf *H* (*t*_*i*_) must satisfy the following condition:

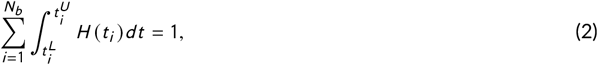

where 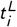 and 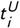 represent the starting and end times for branch *t*_*i*_, respectively. By estimating *λ*_0_ and *H* (*t*_*i*_), it becomes possible to reconstruct the rate of interest, *λ* (*t*_*i*_).

#### Estimating the probability density function *H* (*t*_*i*_)

In standard settings, the function *G* (*t*) is often modeled using a mixture distribution, allowing the observed data to influence the shape of *G* (*t*), which in turn determines how the rate *λ* (*t*) varies over time (Ross and Markwick, 2018). However, specifying a probability measure over the entire timetree is challenging due to the presence of branch splits. To address this, we employ kernel density estimation (KDE) for *H* (*t*_*i*_). KDE is a non-parametric method for estimating pdf.

To ensure that *H* (*t*_*i*_) is a smooth function, particularly over branch split events, we introduce a specialized Markov kernel (MK) for KDE. This kernel conditions the density at a given time point based only on past data, following the plausible evolutionary assumption that present events depend solely on the past. Thus, the KDE estimate 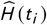 for our pdf can be expressed as:

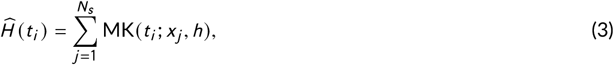

where *x*_*j*_ represents the time point for a state change event, and *h* denotes the smoothing parameter known as the bandwidth. The Markov kernel is a truncated Normal distribution (TN), with the truncated left half (corresponding to the past time). The parameters of MK and TN have the following mapping:

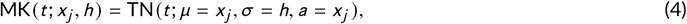

where *µ* represents the mean, *σ* represents the standard deviation, and *a* denotes the lower bound of the truncation. The upper bound of the truncation is undefined (Mersmann et al., 2023).

The Markov kernel, has only one free parameter, the bandwidth *h* to be estimated from the data prior to performing KDE. The choice of bandwidth in KDE is crucial as it affects the trade-off between bias and variance in density estimation. `Ontophylo` offers a range of default R functions (e.g., bw.nrd, bw.ucv, etc) for bandwidth selection (R Core Team, 2023).

The main steps for density estimation in `Ontophylo` are as follows. For a sample of *N*_*sm*_ stochastic maps corresponding to an amalgamated character, the state change events from each map are pooled together onto a single phylogenetic tree. Each branch of the pooled tree is represented by the entire chain of branches from the root to the given branch inclusively. This representation improves the power of kernel density estimation (KDE) at the beginning of each branch. Next, for each branch, the pdf 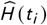 is estimated using Equation (3). Subsequently, the estimated 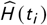 is smoothed using the LOESS method from the fANCOVA package (Wang, 2020) to enhance its smoothness. The final pdf 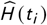 is obtained by integrating it over the entire tree and normalizing it to satisfy Equation (2).

#### Estimating amplitude *λ*_0_

The amplitude parameter *λ*_0_ is estimated using Maximum Likelihood (ML), for which an analytical solution is available. Given a sample of *N*_*sm*_ stochastic maps, the total number of state change events *s*_*i*_ is calculated for each map, where *s*_*i*_ ∈ *s*_1_, *s*_2_, …, *s*_*N*_*sm* . The ML estimate is:

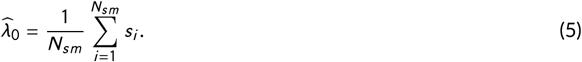

Our package also provides Bayesian inference for *λ*_0_, for which an analytical solution is available too. The conjugate prior for *λ*_0_ is the gamma distribution (Ross and Markwick, 2018). Assuming that *λ*_0_ ∼ Γ(*α, β*), gives the following solution for the posterior:

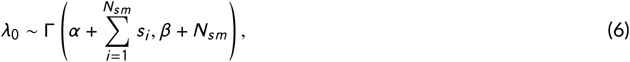

where gamma density is parameterized in terms of shape parameter (*α*) and an inverse scale parameter (*β*). It is worth noting that the posterior mean of *λ*_0_ approaches the ML estimate 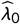 as *α* → 0 and *β* → 0. Therefore, the gamma distribution with low values of *α* and *β* can be used as an uninformative prior for *λ*_0_.

#### Estimating *λ* (*t*_*i*_)

Finally, our parameter of interest, the branch-specific rate *λ* (*t*_*i*_), can be estimated using Equation (1), which involves the product of *λ*_0_ and *H* (*t*_*i*_).

### 2.3 Benchmarking

To evaluate the package performance in estimating rates of discrete character evolution, two sets of simulations were performed. The first set (set 1), aimed to evaluate overall character rates. A tree with 250 taxa was simulated under a bird-death process using geiger (Pennell et al., 2014) resulting in a root age of 117.2 Myr. Two subanalyses were performed.

The first subanalysis (set 1A) assessed the rates for individual characters. 100 characters with two, three, and four states were simulated onto this tree using phytools, resulting in 300 characters in total. For each character simulation, instantaneous rates were sampled from a normal distribution (`mean = 0.5`, `var = 0.05`), and the Q matrix was scaled so as to result in a given number of expected changes per branch length unit. The scaling was done for a scalar of 0.01, 0.1, 1.0, and 10.0, thus resulting in 1200 individual character simulations. For each simulation, 100 stochastic character histories were sampled using `corHMM`, and then overall rates calculated from `corHMM` were compared to the mean rates estimated using `Ontophylo` with the tree discretization parameter set to 500.

The second subanalysis (set 1B) assessed the rates for individual versus amalgamated characters. In this subanalysis, 100 rounds of resampling were performed on 2-state characters from the previous subanalysis. In each round, N characters were sampled, and then the mean of individual character rates (i.e. before amalgamation) estimated with `Ontophylo` was compared to that of the respective amalgamated character. Resampling was done for an N of two, three, and four characters and scalars of 0.1 and 1.0.

The second set of simulations (set 2), aimed to evaluate overall trends over time across the phylogeny in character rates estimated on tree branches. A tree with 100 taxa was simulated as described above resulting in a root age of 71.4 Myr. Binary characters were simulated under three conditions: constant rates, with an 100x decrease, and an x100 increase in rates through time. Simulations were performed for a single character, five amalgamated characters, and ten amalgamated characters. In all cases, the initial root state was set to ‘0’ with a probability of 1, 500 stochastic character histories were sampled using `phytools`, and the tree discretization parameter used by `Ontophylo` was set to 4000.

The R scripts used in the simulations are available in the online Supporting Information (Appendix S1).

### 2.4 Showcase analyses

To showcase the package functionality two data sets were simulated on the Hymenoptera phylogeny modified from Klopfstein et al. (2013). The tree was dated using penalized likelihood as implemented in TreePL (Smith and O’Meara, 2012). The first data set comprised 30 binary characters simulated under equal rates split into three subsets of 10 characters each to emulate different anatomical regions: ‘head’, ‘mesosoma’, and ‘metasoma’. The head characters were simulated with a 100x increase in rates (from a scalar of 0.001 to 0.1) in clade A; mesosoma with a 100x decrease in clade B; and metasoma with a 100x increase in clades C and D (see the tree in the online Supporting Information: Appendix S3). Individual characters were annotated manually with arbitrary terms from the Hymenoptera Anatomy Ontology (HAO) (Yoder et al., 2010) representing the three anatomical regions to allow demonstration of ontology-informed amalgamation (see the table in the online Supporting Information: Appendix S3). This data set was used to show `Ontophylo` Applications 1 and 3 (Figure 1c,e).

The second data set comprised 100 characters simulated to emulate a small phylogenetic character matrix representing the entire ‘phenome’ of a hypothetical hymenopteran insect. 50 binary and 25 multistate parsimony-informative synapomorphic characters were simulated by randomly sampling a node and then attributing a derived state (‘1’ or ‘2’) to the tips; the remaining tips were all assigned the ancestral state ‘0’. The remaining 25 characters were simulated as binary characters under a standard Markov model with equal rates and a scalar of 0.1. The procedure described above aimed to simulate a data set with a high phylogenetic signal so the resulting morphospace is structured. This data set was used to show `Ontophylo` Application 2 (Figure 1d).

For both data sets, characters were simulated using `phytools` with the initial root state set to ‘0’ with a probability of 1; model fitting and stochastic character mapping used `corHMM`; and tree discretization parameter was set to 4000 and 500 respectively. The R scripts used for the example applications are available in the online Supporting Information (Appendix S2).

## 3 RESULTS AND DISCUSSION

### 3.1 Validation

Overall rates of discrete character evolution are correctly estimated with `Ontophylo` under most simulation conditions. Rates for individual characters (simulation set 1A) are congruent with simulated rates within the range between 0.1 and 1.0 for all simulations with two, three, and four states (Figure 2a-b; see the figure for three states in the Appendix S3). Very low rates (0.001) are correctly estimated only for 2-state simulations while very high rates (10.0) are poorly estimated in all cases, likely due to saturation effects. For amalgamated characters (simulation set 1B), rates are correctly estimated for simulated low rates (0.1) (Figure 2c) and always underestimated for simulated high rates (1.0) (Figure 2d). Note that, for simulated low rates, as expected, rates for the 2-, 3- and 4-state amalgamated characters are respectively two, three, and four times the mean rates of their individual characters (Figure 2c: dotted grey lines). Branch rates (simulation set 2) are also correctly estimated under all simulation conditions with one, five, and ten amalgamated characters (Figure 3: dotted red lines): with no overall trends through time (Figure 3a,d,g); decreasing rates through time (Figure 3b,e,h); and increasing rates through time (Figure 3c,f,i). In conjunction with the example applications below, these simulations demonstrate that `Ontophylo` is capable of correctly recovering underlying evolutionary patterns from anatomical data.

**FIGURE 2.**
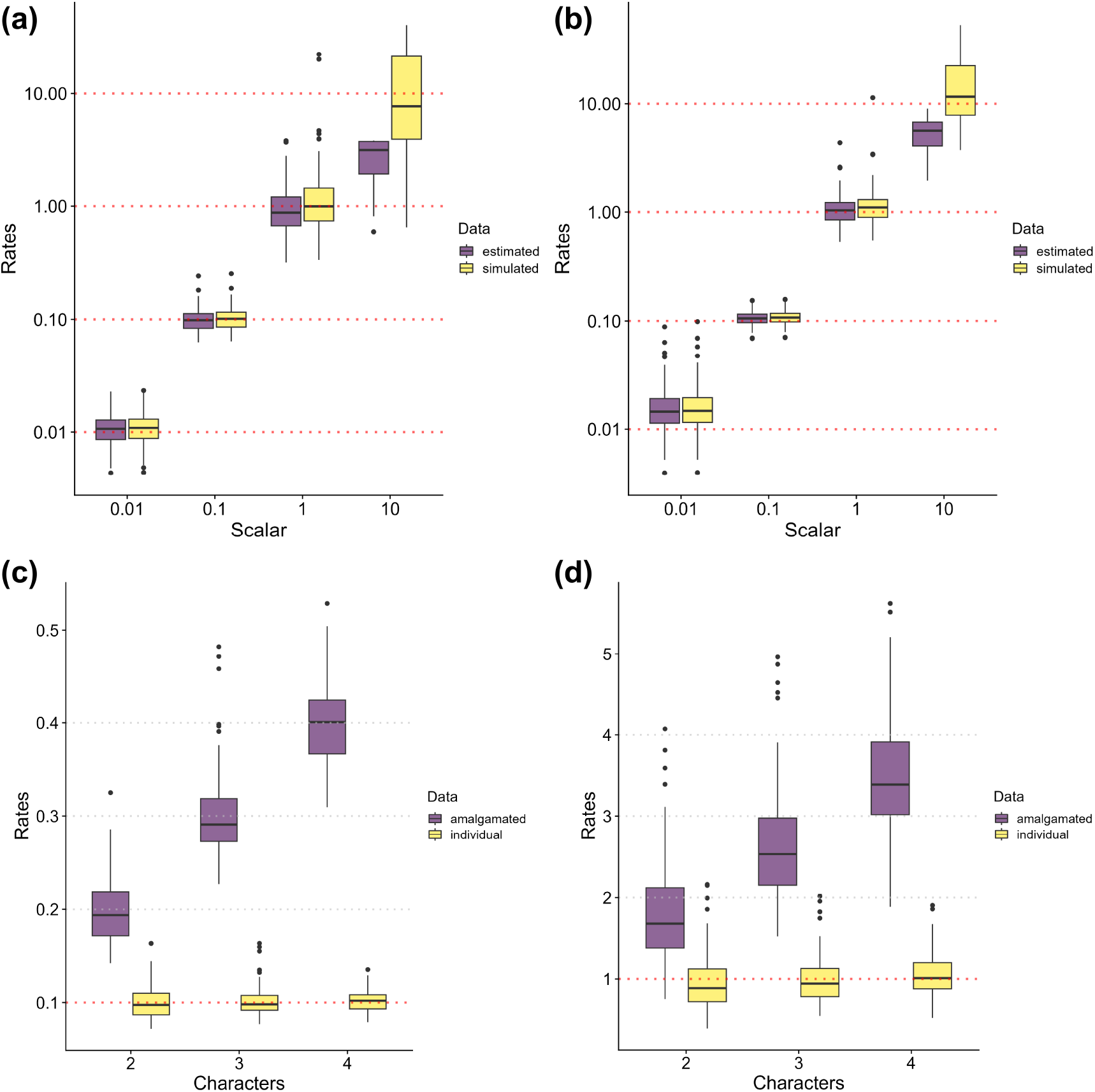
Evaluation of overall rates estimated with *`Ontophylo`*. **(a)** Simulated and estimated rates for 2-state characters. **(b)** Same as (a), but for 4-state characters. **(c)** Estimated rates for amalgamated characters versus the mean rate of corresponding individual characters for simulations under a 0.1 rate scalar. **(d)** Same as (c), but for 1.0 rate scalar. The dotted red line in c-d indicates the rate values for individual characters and the grey line those for amalgamated characters.

**FIGURE 3.**
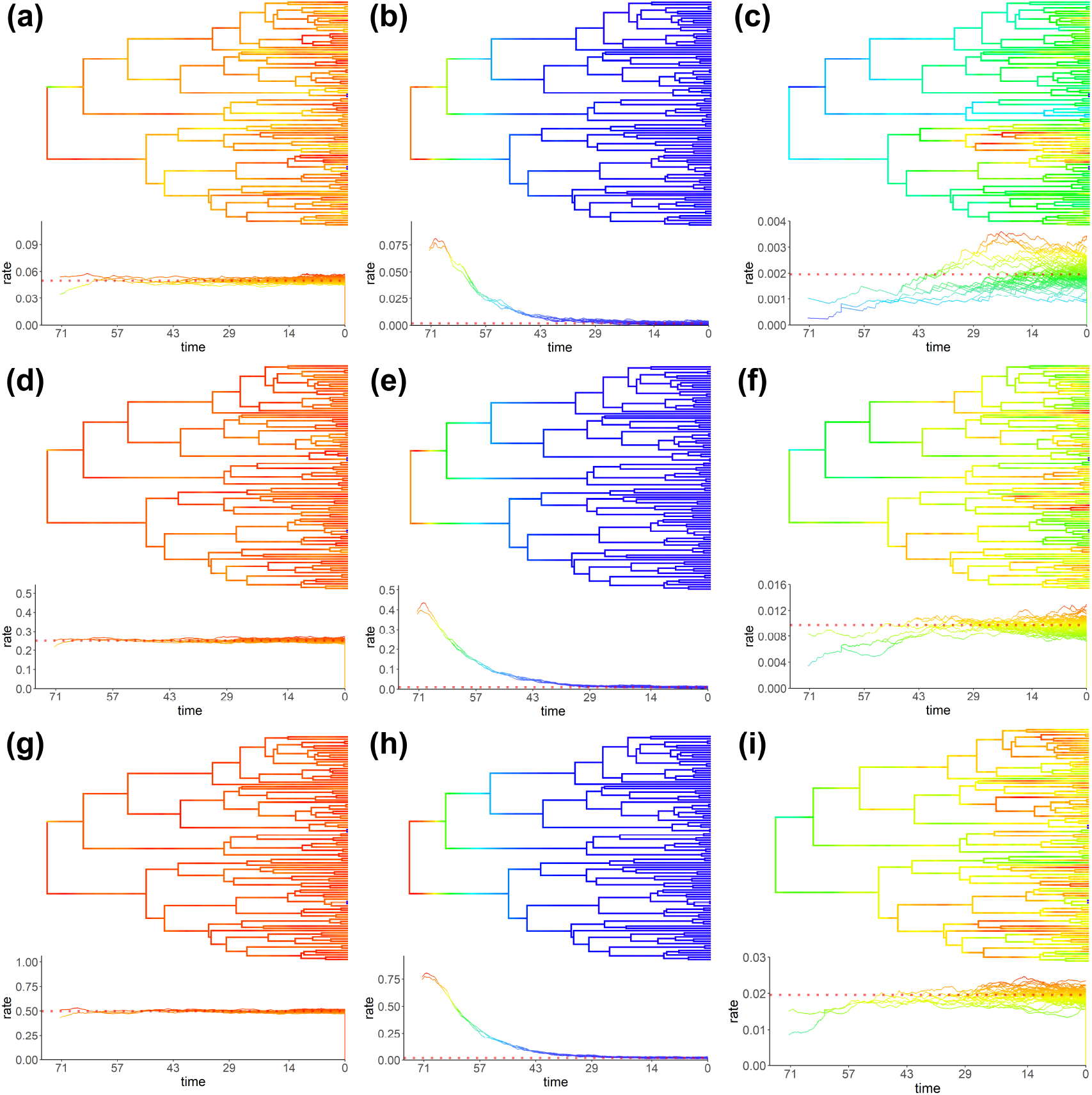
Evaluation of branch rates estimated with *`Ontophylo`*. **(a-c)** A single binary character. **(d-f)** Five binary amalgamated characters. **(g-i)** Ten binary amalgamated characters. Simulations with no rate bias (a,d,g), decreasing rates (b,e,h), and increasing rates (c,f,i) through time. The dotted red line indicates the overall rate value of simulations.

### 3.2 Applications

As an example of common user-case scenarios, the analyses of simulated Hymenoptera data sets show the utility of employing `Ontophylo` in comparative studies of anatomical data. Evolutionary rates of traits from different anatomical entities likely vary across characters, morphological complexes, anatomical regions, or the entire phenome (Wagner, 2012). Furthermore, rate heterogeneity across lineages is usually expected for old and diverse clades across the Tree of Life (Beaulieu et al., 2013). When rate variation occurs—across anatomy or phylogeny—`Ontophylo` helps to unveil the tempo and mode of morphological evolution.

By assessing the mean branch rates estimated for different anatomical regions and the entire phenome a researcher can investigate clade-specific patterns and temporal trends of morphological evolution. In the analyses of the simulated Hymenoptera data set, an increase in rates of head characters is correctly recovered for clade A around 225 Myr (Figure 4a); a decrease in rates of mesosoma characters for clade B around 275 Myr (Figure 4b); and an increase in rates of metasoma characters for clade C around 175 Myr and to a lesser degree for clade D around 90 Myr (Figure 4c). For the entire phenome, the three peaks in rates are congruent with the ones expected from the individual anatomical regions (Figure 4d).

**FIGURE 4.**
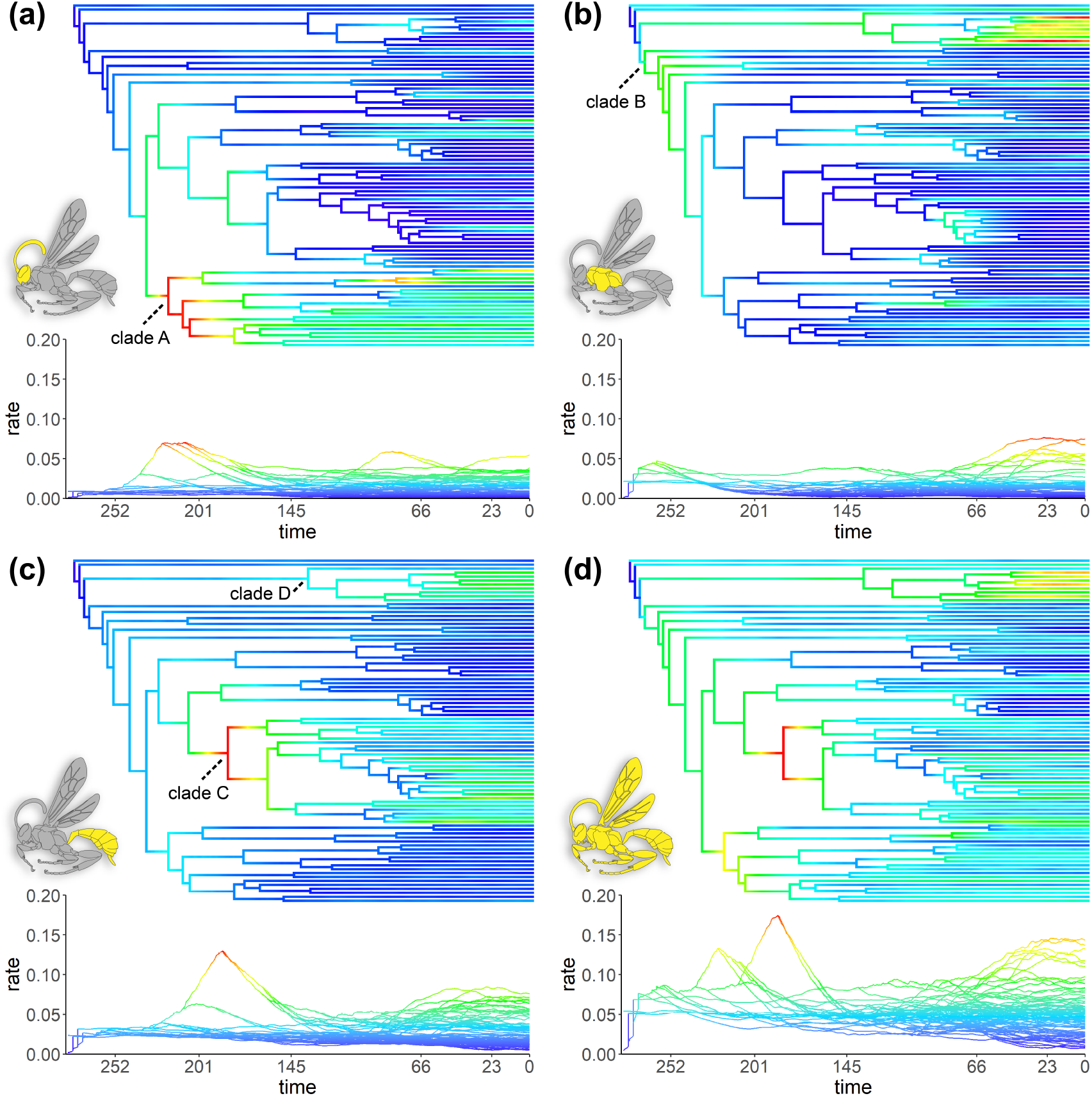
Contmap of mean branch rates and edgeplots for three anatomical regions and phenome for the first simulated example data set of Hymenoptera. **(a)** Head. **(b)** Mesosoma. **(c)** Metasoma. **(d)** Phenome. Clades A-D represent simulated shifts in rates as explained in the materials and methods.

A researcher can also investigate rate variation across anatomical regions. For example, in the simulated Hymenoptera data set, low rates are recovered for all anatomical regions at branch 1; slightly higher for head and metasoma at branch 2; much higher for metasoma at branch 3; and lower again for all anatomical regions at branch 4 (Figure 5).

**FIGURE 5.**
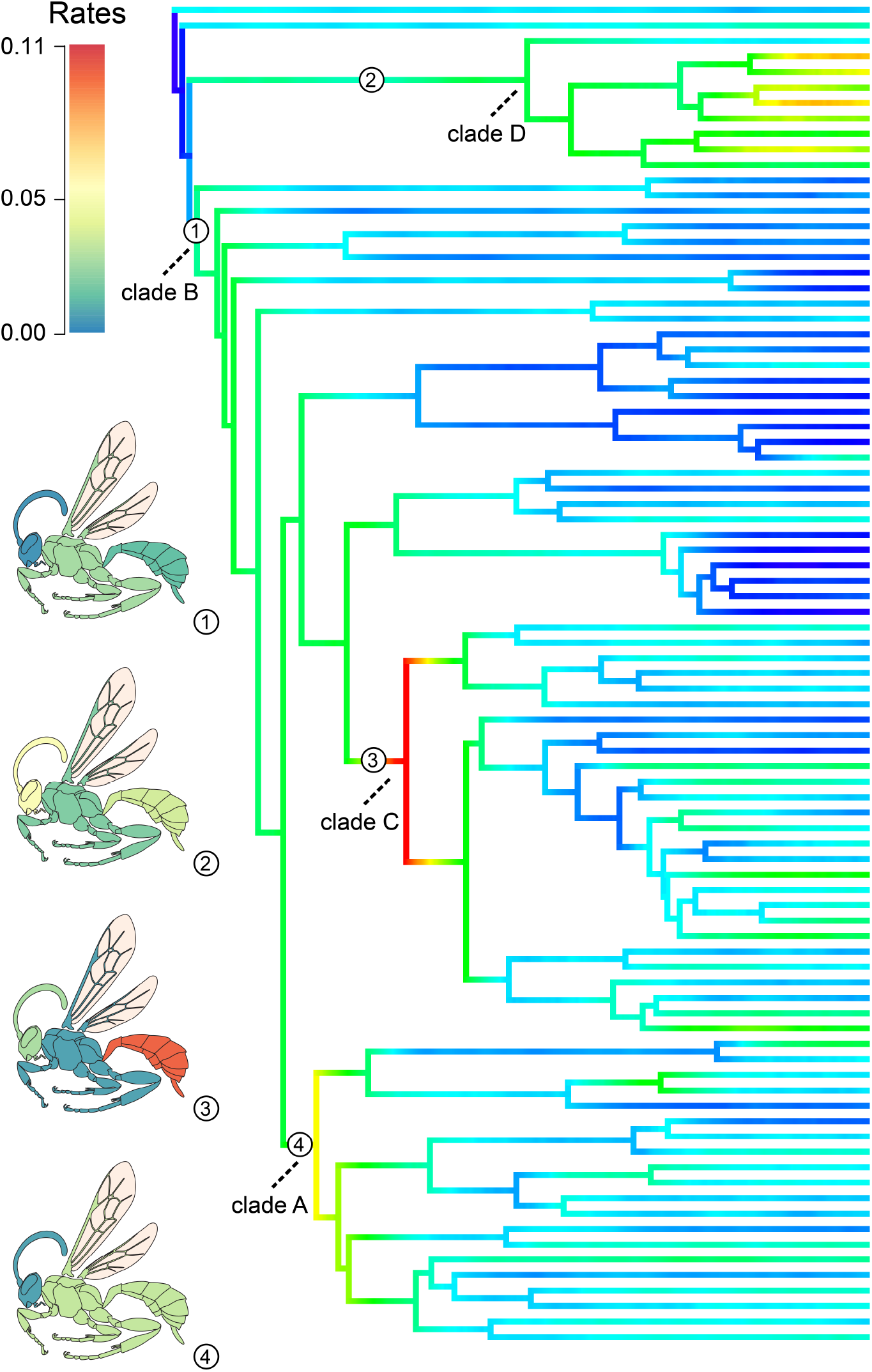
Evolutionary rates for three anatomical regions (head, mesosoma, and metasoma) at different selected branches (1-4) of the phenome contmap. Clades A-D represent simulated shifts in rates as explained in the materials and methods.

Another advantage of employing `Ontophylo` is the possibility of investigating how morphospace occupation changes through time since amalgamated character state reconstructions are available for all tree branches at multiple time slices. For example, in the simulated Hymenoptera data set the phenome morphospace occupation starts around 280 Myr (Figure 6a: purple dot), proceeds gradually as early branches are visited and new phenomic states are sampled around 230 Myr (Figure 6b: purple dots), and finally spreads towards three different regions of the morphospace, roughly corresponding to early diverging symphytan lineages and two main clades in Apocrita from around 130 Myr (Figure 6c: purple to grey dots) to the present (Figure 6d: grey to green dots).

**FIGURE 6.**
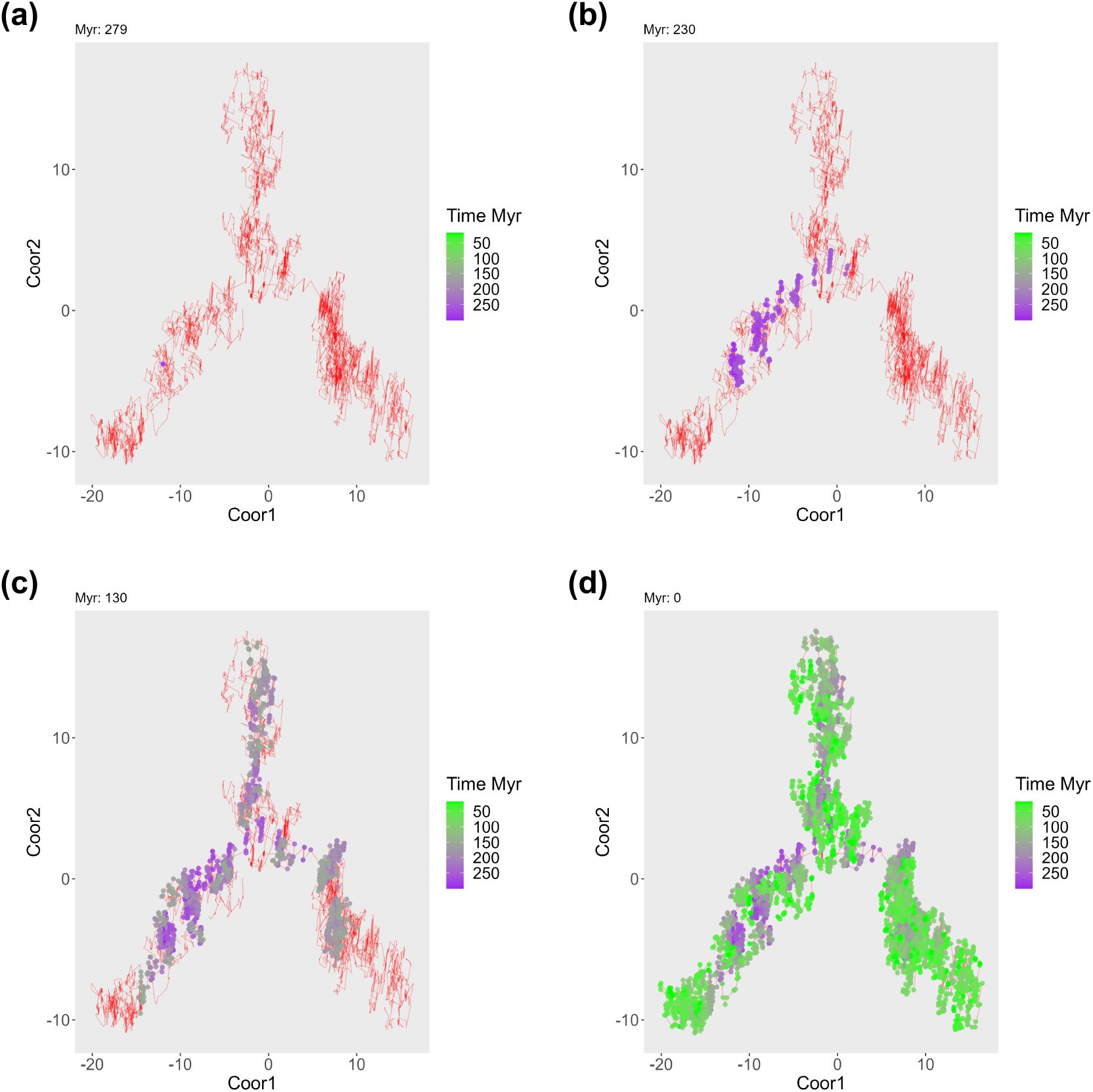
Morphospace reconstruction for the simulated phenome of Hymenoptera. The red lines in **(a-b)** represent tree branches plotted in 2D after applying a dimension reduction technique (in this case MDS); a SM explicitly reconstructs ancestral states for every point in a tree, and MDS was applied to those states; the obtained 2D plot indicates similarities between the branches of the tree based on Hamming distances between the reconstructed states. The evolutionary rate over time is shown by using points whose colors indicate the time when the states appeared. This figure illustrates how the morphospace becomes filled for four time slices. **(a)** 279 Myr. **(b)** 279-230 Myr. **(c)** 279-130 Myr. **(d)** 279-present. A morphospace animation is available in Supporting Information: Appendix S4

The cases presented above show just some examples of what can be potentially investigated with `Ontophylo`. Other stimulating evolutionary questions can be also further addressed. How variable is the rate of evolution of the entire phenome across lineages and over timefl Which scenario best describes the evolution of phenomes over time—constant rate, early burst, acceleration toward the present, or something elsefl Do the evolutionary rates of different body regions exhibit a common trend, or are their evolution uniquefl In what ways does morphospace expand and become occupied over timefl `Ontophylo` can be used to address questions on rate correlation between body regions and other biotic and abiotic variables by providing estimates of tip rates that can be employed for standard phylogenetic regression methods.

### 3.3 Practical recommendations

For many functions in the workflow of `Ontophylo`, two parameters are key: resolution and bandwidth. The resolution controls the size of the episodic bins used to discretize tree branches. Discretization is important to allow amalgamation by stacking stochastic maps from individual characters. Higher resolution results in more bins with a smaller size. The number of bins will directly impact KDE estimation and thus, rates estimated from pNHPP. Although we recommend using as higher a value of resolution as possible, there is a trade-off between resolution and computational eflciency. In most cases, a resolution of around 500 should be enough to give a good fine-tuning of the analyses. The bandwidth controls the smoothing of the KDE estimation. The choice of bandwidth in KDE is crucial as it affects the trade-off between bias and variance in density estimation. A small bandwidth can lead to overfitting, where the density estimate becomes too sensitive to individual data points, resulting in a spiky and noisy estimate. On the other hand, a large bandwidth can cause oversmoothing, where the estimate becomes too generalized, potentially obscuring important features of the data. There are various methods available to select an appropriate bandwidth for KDE, such as cross-validation, plug-in methods, or rule-of-thumb approaches like Silverman’s rule (R Core Team, 2023; Silverman, 1986). These methods aim to strike a balance between capturing the structure of the data while avoiding excessive bias or variance in the density estimate. Currently, `Ontophylo` uses standard bandwidth selectors from R stats, and we recommend trying different options to evaluate the effects on the smoothing.

### 3.4 Limitations

To harness the full potential of `Ontophylo`, phylogenetic characters should be annotated with ontology terms to allow character amalgamation at multiple levels. The annotations do not necessarily need to comprise full semantic statements such as those expressed with the entity-quality model (i.e. one term from an anatomy ontology combined with another from a quality ontology) (Balhoff et al., 2014). Only terms referring to anatomical entities are needed. Annotations can be done semi-automatically with ontoFAST (Tarasov et al., 2022), but in most cases, the final decision on the selection of terms to be used depends on expert judgment.

Dependencies among characters from different anatomical entities should be dealt with before moving to the amalgamation step. Automatic setup of evolutionary models used for stochastic character mapping can be done with rphenoscate (Porto et al., 2023) using the ontological knowledge on anatomical dependencies, but there still is a discussion in the literature on how to properly translate this knowledge into models and coding schemes (Tarasov, 2023; Simões et al., 2023; Tarasov, 2019).

Although `Ontophylo` performs relatively well for data sets within the range of 100 to 200 taxa and 100 amalgamated characters, we expect that some current applications (such as the morphospace dynamics reconstruction) might not scale well as the size of data sets increases drastically. That is especially true when the resolution parameter is high.

Finally, one obvious factor that can influence the rates estimated with `Ontophylo` is the bias in character selection. The rates for the amalgamated character will be heavily influenced by the number of individual characters and the number of states of each character. With more characters and states, assuming characters are independent, a higher number of transitions is to be expected. With this in mind, researchers should be aware that comparisons between rates estimated for amalgamated characters representing different anatomical regions will usually be biased toward anatomical regions with more individual characters. Nonetheless, comparisons for the same body region across different branches of the tree are still meaningful.

### 3.5 Phenomics

In the era of phenomic-scale data sets in evolutionary biology, taxonomic literature is a vast and rich source of knowledge, usually available as photographs, illustrations, species descriptions, and phylogenetic character matrices (Deans et al., 2012, 2015; Thessen et al., 2020). Such knowledge can be made accessible to computer reasoning by annotating images and statements about organismal anatomy with terms from ontologies. We envision that `Ontophylo` can be integrated with other packages such as ontoFAST (Tarasov et al., 2022), ontobayes (Porto et al., 2022), `rphenoscape` (https://github.com/phenoscape/rphenoscape), and rphenoscate (Porto et al., 2023) as part of a growing ecosystem of R packages integrating ontologies and phylogenetics. There are already tools available for image segmentation (Lösel et al., 2020; Schwartz and Alfaro, 2021), annotating phylogenetic characters (Balhoff et al., 2010, 2014), and writing semantic descriptions (Balhoff et al., 2013; Mikó et al., 2021). By integrating semantic data, ontologies, and phylogenetics, new opportunities are rapidly emerging to investigate questions about organismal anatomy at an unprecedented scale taking advantage of the fields of computer vision, deep learning, and robotics applied to taxonomy, ecology, and evolution (Lürig et al., 2021; Borowiec et al., 2022; Wührl et al., 2022).

Furthermore, `Ontophylo` applications can be expanded and complemented in three different ways: (1) by adding alternative tools to assess and visualize rate magnitudes and rate shifts on branches; (2) by incorporating better ways to assess and visualize uncertainty and variance of rate estimates; (3) by adding alternative methods of dimensionality reduction other than multidimensional scaling (MDS) and other metrics to compare phenotypes other than Hamming distances, the current defaults of Application 2.

### 3.6 Conclusions

In this study, we have demonstrated how `Ontophylo` can be used to study the tempo and mode of phenome evolution by capitalizing on current advances in phylogenetics and integration with ontological knowledge. We hope our new package will prompt researchers to ask new questions about phenome evolution in different groups of organisms and to start a movement to annotate legacy anatomical and phylogenetic data with ontology terms and produce new semantically-enriched data.

## 4 ACKNOWLEDGEMENTS

This work received funding from the National Science Foundation (NSF 1661516 to J.C.U.) and the Academy of Finland (346294 and 339576 to S.T.).

## 6 AUTHOR CONTRIBUTIONS

ST conceived the package; DSP and ST designed, tested, and wrote the documentation of the package; all authors contributed to the draft of the manuscript and revised the final version of the paper.

## 7 DATA AVAILABILITY STATEMENT

The current development version of `Ontophylo` is available on GitHub (https://github.com/diegosasso/Ontophylo). The code of version 1.0.0 and all data and scripts of simulations are archived on Zenodo (XXXX).

## 9 SUPPORTING INFORMATION

Additional supporting information can be found online in the Supporting Information section at the end of this article.

**Appendix S1:** R code for the benchmarking simulations.

**Appendix S2:** R code for the example applications. **Appendix S3:** Supplementary tables and figures. **Appendix S4:** Morphospace animation for Figure 6.

